# Membrane interactions and uncoating of Aichi virus, a picornavirus that lacks a VP4

**DOI:** 10.1101/2022.01.10.475380

**Authors:** James T. Kelly, Jessica Swanson, Joseph Newman, Elisabetta Groppelli, Nicola J. Stonehouse, Tobias J. Tuthill

**Affiliations:** The Pirbright Institute, Ash Road, Pirbright, GU24 0NF, UK; School of Molecular and Cellular Biology, Faculty of Biological Sciences and Astbury Centre for Structural Molecular Biology, University of Leeds, Leeds, LS2 9JT, UK; Institute for Infection and Immunity, St. George’s University of London, Tooting, London, SW17 0RE, UK

**Keywords:** Aichi virus, Kobuvirus, Picornavirus, VP4, VP0, uncoating, pore formation, viroporin

## Abstract

Kobuviruses are an unusual and poorly characterised genus within the picornavirus family, and can cause gastrointestinal enteric disease in humans, livestock and pets. The human Kobuvirus, Aichi virus (AiV) can cause severe gastroenteritis and deaths in children below the age of five years, however this is a very rare occurrence. During the assembly of most picornaviruses (e.g. poliovirus, rhinovirus and foot-and-mouth disease virus), the capsid precursor protein VP0 is cleaved into VP4 and VP2. However, Kobuviruses retain an uncleaved VP0. From studies with other picornaviruses, it is known that VP4 performs the essential function of pore formation in membranes, which facilitates transfer of the viral genome across the endosomal membrane and into the cytoplasm for replication. Here, we employ genome exposure and membrane interaction assays to demonstrate that pH plays a critical role in AiV uncoating and membrane interactions. We demonstrate that incubation at low pH alters the exposure of hydrophobic residues within the capsid, enhances genome exposure and enhances permeabilisation of model membranes. Furthermore, using peptides we demonstrate that the N-terminus of VP0 mediates membrane pore formation in model membranes, indicating that this plays an analogous function to VP4.

**Importance:** To initiate infection, viruses must enter a host cell and deliver their genome into the appropriate location. The picornavirus family of small non-enveloped RNA viruses includes significant human and animal pathogens and are also models to understand the process of cell entry. Most picornavirus capsids contain the internal protein VP4, generated from cleavage of a VP0 precursor. During entry, VP4 is released from the capsid. In enteroviruses this forms a membrane pore, which facilitates genome release into the cytoplasm. Due to high levels of sequence similarity, it is expected to play the same role for other picornaviruses. Some picornaviruses, such as Aichi virus, retain an intact VP0, and it is unknown how these viruses re-arrange their capsids and induce membrane permeability in the absence of VP4. Here we have used Aichi virus as a model VP0 virus to test for conservation of function between VP0 and VP4. This could enhance understanding of pore function and lead to development of novel therapeutic agents that block entry.

## Introduction

For many non-enveloped viruses, replication depends on the capsid first binding a receptor to trigger uptake into a cell via endocytosis. During entry the virus must deliver its RNA genome into the cytoplasm. Mechanisms of delivery are not well understood, but the proposed mechanism in picornaviruses (such as poliovirus (PV) and human rhinoviruses (RV), involves capsid structural rearrangements that enable the virus to interact with the endosomal membrane and form a pore. The capsid then uncoats, releasing its genome through the pore, across the endosomal membrane and into the cytoplasm. In many picornaviruses and picorna-like viruses, viral capsid protein VP4 is a small internal component of the capsid that is released during cell entry to initiate pore formation (1–7). VP4 is formed from the cleavage of capsid protein VP0 into VP2 and VP4 (8–10). However, certain picornavirus do not undergo VP0 cleavage, therefore do not possess a VP4 protein, and it is unknown what component of the capsid performs the normal function of VP4.

The best characterised picornavirus genera that possess uncleaved VP0 are Kobuviruses and parechoviruses. Kobuviruses are associated with cases of acute gastroenteritis in people, livestock and pets (11–14), including the best studied member, the human pathogen Aichi virus (AiV). The virus is wide-spread, with 80% to 95% of adults reportedly having antibodies against the virus (15–17). AiV is generally asymptomatic, however it can cause mild gastrointestinal upset and there have even been fatal cases reported in children under five, especially in developing countries (14, 18–20).

Picornavirus particles consist of a single positive sense RNA genome, within a non-enveloped capsid composed of 60 copies of four structural proteins, VP1, VP2, VP3 and VP4 arranged in pseudo T=3 icosahedral symmetry. In the majority of picornaviruses, VP2 and VP4 are derived from a precursor called VP0, which undergoes a maturation cleavage to form VP2 and VP4 (e.g. enteroviruses, aphthoviruses, cardioviruses and hepatoviruses), this is thought to be triggered by RNA encapsidation in some viruses (8–10, 21). However for the Kobuvirus and parechovirus genera, VP0 does not cleave and the mature capsid contains an intact VP0 (22, 23). In VP4-containing viruses, VP4 is usually myristoylated and by using specific inhibitors or mutagenesis of a myristoylation signal sequence to prevent myristoylation, it has been shown to play a critical role in virus assembly and entry (24–27). The N-terminus of Kobuvirus VP0, but not parechovirus VP0 is myristoylated (22, 28), however, myristoylation seems unlikely to play an essential role as the specific inhibitors are unable to restrict infection by these viruses (28). Uncoating has been extensively studied in VP4-containing picornaviruses, especially enteroviruses (e.g. PV, RV). However, there are few studies on uncoating in VP0 containing viruses and no studies have been reported on the role of VP0 in uncoating, although it is assumed that the N-terminus of VP0 may be involved in pore formation as this is in an analogous location to VP4.

In order to uncoat and form a pore within the endosomal membrane, viral capsids must undergo extensive structural rearrangements. Experimental and structural studies of different types of picornavirus particles have given great insights into the structural rearrangements that occur during uncoating of VP4-containing picornaviruses (29–37). The trigger varies between viruses, for aphthoviruses these changes can be initiated solely by exposure to low pH (38, 39) while for major group RV receptor interactions in combination with endosomal acidification are required (40–43). AiV capsids are known to be destabilised by low pH, and therefore endosomal acidification may play a role in AiV uncoating (44).

Studies with enterovirus particles have shown that these viruses are able to bind to and permeabilise membranes, during a process known as capsid breathing (2, 45). Studies using intact virions, peptides of VP1-N and VP4 and antibodies raised against VP1-N and VP4, in conjunction with membrane binding and pore formation assays revealed the N-terminus of VP1 is involved in attaching the enterovirus capsid to the membrane and VP4 is involved in pore formation (7, 45–47). We have shown that recombinant VP4 and VP4-peptides of rhinovirus 16 (RV16) form size selective pores in model membranes consistent with the size of a single strand of RNA to pass thorough (2, 7). Furthermore, mutation of residue T28 in PV VP4 can reduce the capsids ability to permeabilise model membranes (5). In combination with this biophysical data, structural studies have helped develop a model for enterovirus uncoating. Incubation of enterovirus particles with their receptor or heating past physiological temperature, can trigger mature particles to uncoat into uncoating intermediate particles (Altered particles/A-particles) or empty particles (30). A-particles still contain the viral genome, but the capsid has undergone expansion and structural rearrangements, including VP4 release and externalisation of N-terminus of VP1, whereas empty particles have released their genome and undergone further structural rearrangements (29–37). Biophysical and structural data of PV in the presence of model membranes indicate that VP4 and the N-terminus of VP1 together may form an ‘umbilicus’ which tethers the virus to the endosomal membrane (46, 48).

Unlike enteroviruses, the aphthoviruses and cardioviruses do not produce stable empty capsids during uncoating *in vitro*. The aphthoviruses foot-and-mouth disease virus (FMDV) and equine rhinitis A virus (ERAV), disassemble into pentamers almost instantly after exposure to a pH critical for uncoating. Exposure to heat also induces disassembly. An uncoating intermediate/empty particle structure has been solved for ERAV from crystals grown at low pH (38). From this it was observed that VP4 is completely released from the capsid and the N-terminus of VP2 may be externalised from the capsid, rather than VP1 as in enteroviruses. Due to this, it is hypothesised that the N-terminus of VP2 will be involved in endosomal tethering in aphthoviruses (38). Similarly, cardioviruses also do not produce stable empty particles, for example, heating of Saffold virus 3 to 42 °C for 5 minutes induces particle disassembly, however exposure for 2 minutes induces a mixture of empty and A-particles (49). A low resolution cryo-EM structure of the A-particles, shows the particles have expanded, released VP4 and that there is an unconnected density leaving the particle that might be the VP1 N-terminal arm (49). To summarise, the capsid rearrangements that occur in VP4-containing picornaviruses, involve capsid expansion, release of VP4 and externalisation of the N-terminus of either VP1 or VP2.

However, less is known about uncoating in VP0-containing picornaviruses. To date AiV is the only VP0-containing picornavirus for which an empty capsid structure has been determined. Cryo-EM structures of mature and empty AiV capsids produced by heating revealed that in the empty particle the N-terminus of VP0 and VP1 become disordered, however no proteins were observed to be released or externalised (50). This differs to what is known about VP4-containing picornaviruses, where VP4 is released and the N-terminus of either VP2 or VP1 becomes externalised. It is difficult to envisage a model for membrane interactions in which capsid proteins are not externalised. For AiV, it was proposed that during uncoating, capsid proteins become externalised but slide back inside the capsid after genome release (50). However, whether this observation is biologically relevant remains to be resolved, as it seems unlikely that capsid proteins could easily be re-internalised if they are inserted within a membrane.

In this study the mechanism of AiV uncoating is analysed using purified virus particles and a VP0 peptide. Using chemical inhibition assays, capsid stability assays and liposome assays with purified virus we show that acidification is essential for AiV entry and that uncoating and membrane interactions are also dependent on low pH. Using peptides in liposome assays we also show that the N-terminus of AiV VP0 plays an equivalent role to that seen in the VP4 proteins of other picornaviruses. Sequence alignments suggest this function may be conserved between all Kobuviruses.

## Results

### AiV endocytosis is dependent on endosome acidification

AiV capsids, in common with those of many other picornaviruses are destabilised by low pH (51), suggesting that endosomal acidification may be a trigger for AiV uncoating. To test if endosome acidification was required for AiV entry, the virus was grown in the presence of the endosome acidification inhibitor NH_4_Cl at a range of non-toxic concentrations. In untreated cultures or those treated with low concentrations (2 mM) of NH_4_Cl, virus infection resulted in complete CPE at 20 hpi, with viral titres of 2-3 × 10^6^ pfu/ml. In contrast, in cultures treated with NH_4_Cl at concentrations of 10 mM and above, the addition of virus did not lead to visible CPE and AiV titres at 20 hpi were reduced by over 99% (Figure 1A)). A time of addition study was then carried out to determine at what point in the virus life cycle NH_4_Cl was inhibitory. An inhibitory concentration of NH_4_Cl was added every hour to a different well of AiV infected cells. This ranged from one hour prior to infection, to 4 hours post infection. This revealed that NH_4_Cl was only inhibitory when added to cells prior to or during infection (Figure 1B). When added 1 hour post infection, little reduction in titre was observed and no reduction occurred when added 2 or more hours post infection (Figure 1B). This shows that NH_4_Cl inhibits AiV infection during entry and not at another part of replication. This is consistent with the requirement for low pH for cell entry.

**Figure 1.**
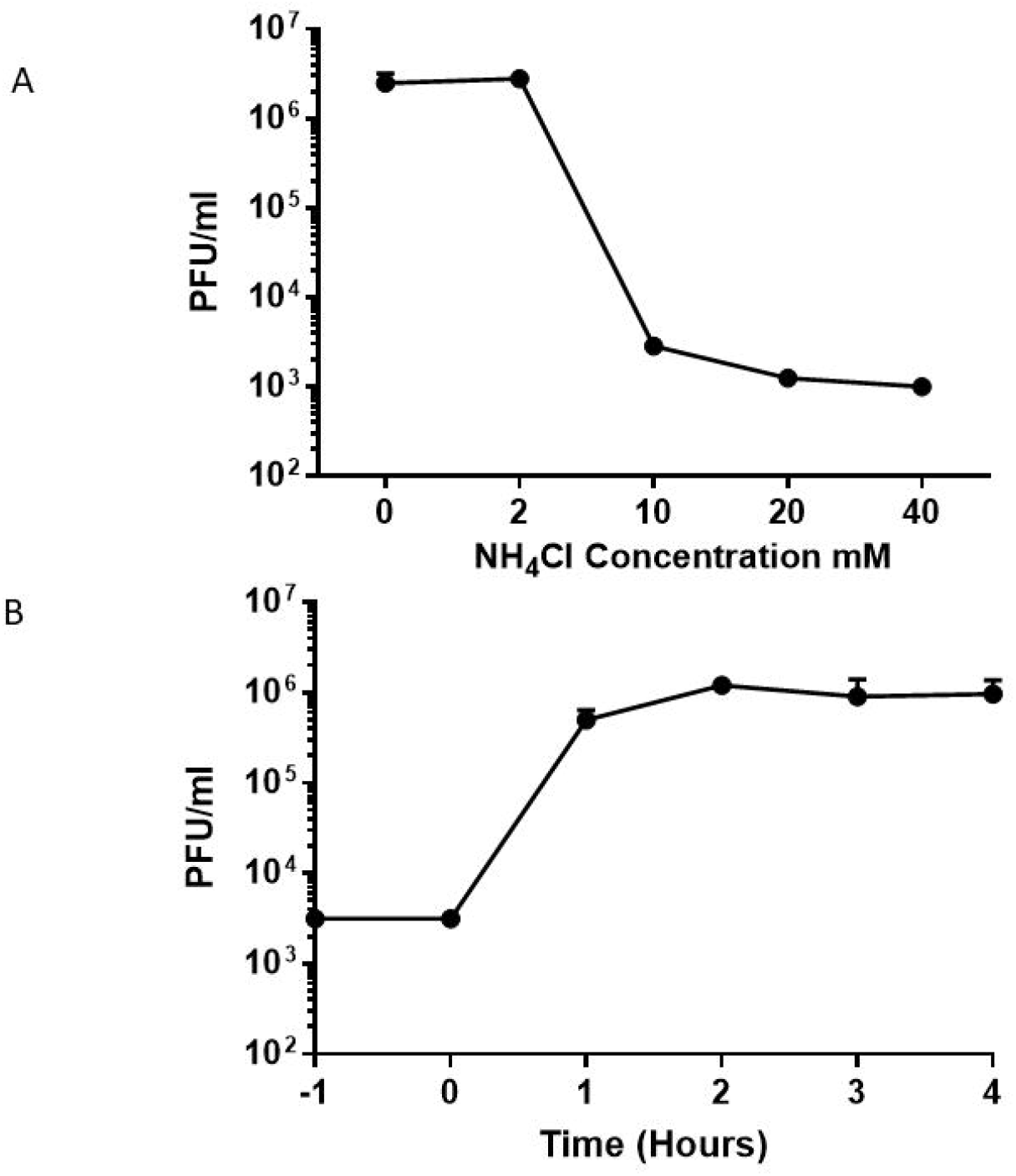
Inhibition of endosomal acidification interferes with an early step in the AiV life cycle. a) Growth of AiV is prevented by treatment of cells with NH_4_Cl: Titre of virus 24 hours post infection of cells treated without, or with 2 mM, 10 mM, 20 mM or 40 mM NH_4_Cl for 2 hours before infection with AiV at MOI 1. b) Time of addition of NH_4_Cl shows effect early in infection: Titre of virus 24 hours post infection of cells treated with 40 mM NH_4_Cl 1 hour before infection (−1) or 0, 1, 2, 3, 4 hours post infection. Experiments were performed in triplicate.

### Decreasing pH enhances capsid alterations

Having shown that endosome acidification is important for AiV entry, we wanted to investigate the effect of pH on genome exposure and capsid protein dynamics. This was assessed using a particle stability thermal release assay (PaSTRy) assay, which has previously been used to study uncoating dynamics and particle stability of enteroviruses, FMDV and AiV (44, 52–55). To perform this, purified virus was incubated at a range of pH values with two dyes (SYTO9 and SYPRO orange). SYTO9 binds and fluoresces in the presence of nucleic acid, indicating genome accessibility. SYPRO orange binds and fluoresces in the presence of hydrophobic amino acid residues, indicating exposure of hydrophobic residues within the capsid proteins. During the assay, temperature was raised by 1 °C every 30 seconds and the level of florescence of each dye was measured.

Previous studies with PaSTRy established that low pH can promote AiV genome exposure to occur at lower temperatures (44). Here we have repeated this study using a finer range of pH values, between pH 7.0 and pH 4.9 (7.0, 6.2, 5.9, 5.6, 5.0 and 4.9), while also tracking the exposure of hydrophobic protein residues. Assays were performed after pre-incubating purified particles for 10 minutes at either room temperature, 56 °C or 59 °C, and were then chilled on ice for 2 minutes, prior to performing the assay.

With a room temperature pre-incubation, SYTO 9 fluorescence began to be detectable between 48 and 49 °C for pH 7.0, 6.2 and 5.9 with maximal fluorescence occurring at 55 °C. At pH 5.6 these values were reduced to 45 and 54 °C respectively. At pH 5 the fluorescence started at 42 °C peaking at 51 °C and at pH 4.9 these values were reduced to 41 °C and 50 °C respectively (Figure 2A). This showed that incubation at lower pH reduces the temperature for RNA exposure. When experiments were repeated with samples pre-incubated at 56 °C, a greatly reduced peak signal was observed at pH 7.0 and no peak was observed at pH 5.0 (Figure 2 B). For samples pre-heated to 59 °C no SYTO9 peak was observed for samples (pH 7.0, 6.2, 5.6, 5.0 4.9) (Figure 2C).

**Figure 2.**
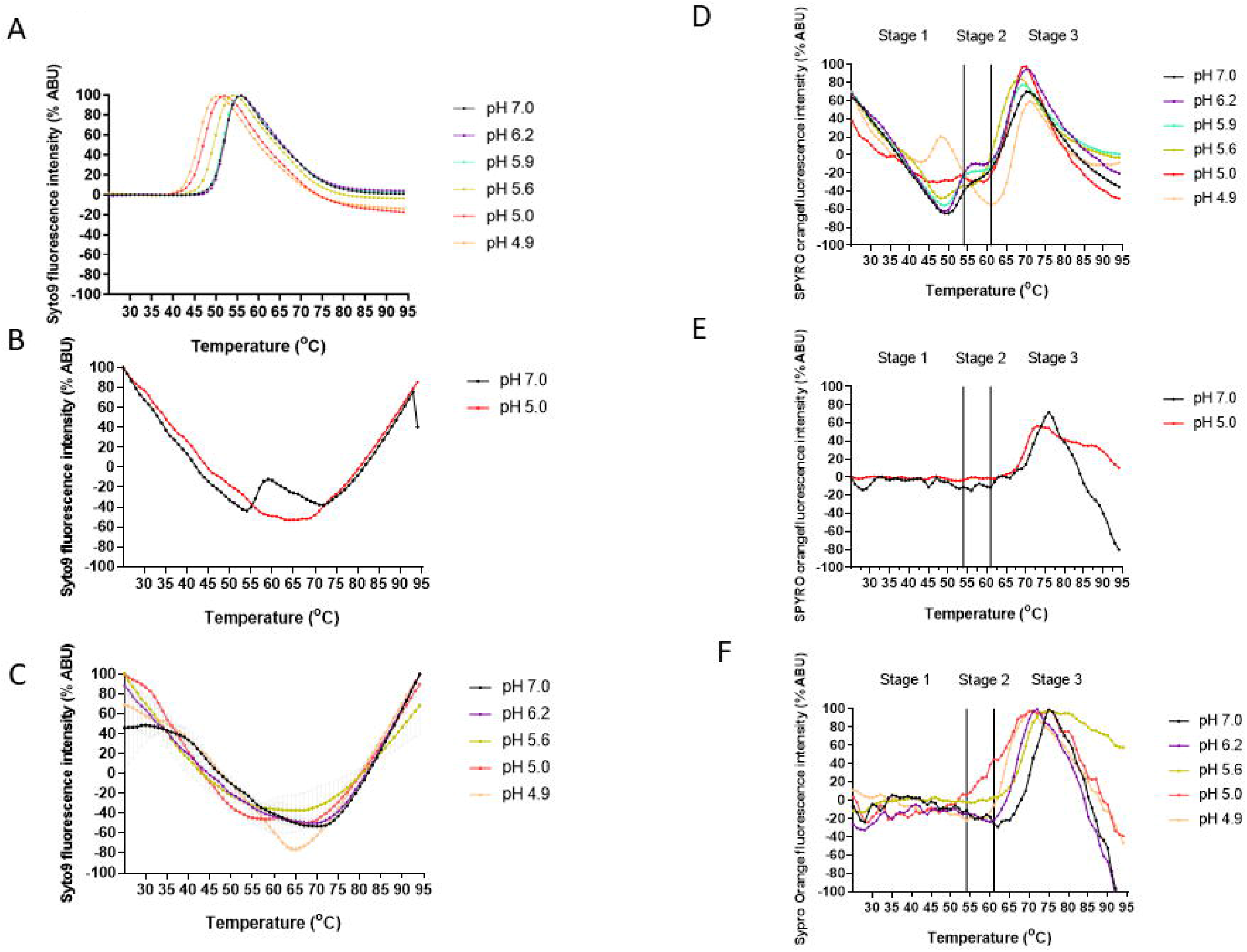
Low pH enhances AiV capsid alterations. AiV analysed by PaSTRy over a range of pH values (pH 7.0, 6.2, 5.9, 5.6, 5.0 and 4.9) with incremental increases of temperature of 1°C every 30 seconds. Experiments were either put in at room temperature (a,d) or heated at either 56 °C (b,e) or 59 °C (c,f) for 10 mins and then chilled on ice for 2 minutes prior to being put on the Stratagene MX3005p quantitative-PCR (qPCR) system. a-c) Relative fluorescence of SYTO9 nucleic acid-binding dye, where increasing signal infers exposure of viral RNA; d-e) Relative fluorescence of SYPRO orange hydrophobic protein residue-binding dye, where increasing signal infers exposure of hydrophobic capsid components. All results are normalised to maximum signal for each experiment, representing 100% signal. All experiments were performed in triplicate, this is a representative trace.

Therefore, overall, as pH decreased, the exposure of nucleic acid appeared to occur at lower temperatures. Results from the pre-incubation at 56 °C indicate that low pH has enhanced genome release and not just exposure, as a peak being present at pH 7.0 but not pH 5.0 indicates that at pH 7.0 some RNA remains within the capsid but at pH 5.0 it has been completely released.

The effect that different pH values had on the profile of hydrophobic protein dynamics as measured by fluorescence is more complex. For simplicity we have separated the profiles into three stages based on events occurring at specific temperatures (Figure 2D). Stage 1 (25-54 °C) for samples pre-incubated at room temperature is characterised by a trough at pH 7.0, 6.2, 5.9 and 5.6, the trough becomes increasingly shallow at lower pH values. At pH 5.0 no trough was observed and instead, the profile resembled a flat line, finally at pH 4.9 a peak was observed. The peak for the hydrophobic protein fluorescence at pH 4.9 occurred 2 °C before the maximum peak of RNA exposure peak (Figure 2B). Stage 2 (54 to 62 °C) is characterised by a sloping shoulder at pH 7, at pH 6.2 and 5.9 the shoulder is flatter, at pH 5.6 and 5 it is almost indistinguishable from the trough in Stage 1 and at pH 4.9 it has become a distinct trough. The beginning of the Stage 2 coincides with maximum RNA exposure for pH 7, 6.2 and 5.9. At Stage 3 (62 to 95 °C) a final large peak is observed in all conditions. The exact temperature that the peak occurs and its magnitude varies between different pH values, this likely represents protein denaturation and is not physiologically relevant. Profiles of AiV at pH 4.9 resemble PaSTRy assay previously observed for enteroviruses at neutral pH, however AiV profiles differ when incubated between pH 7.0 and 5.0 (53, 54). When samples were pre-incubated to 56 or 59 °C a flat line was observed at Stage 1 and Stage 2 but the final Stage 3 peak was still observed (Figure 2 E,F).

The experiments described here have shown that reductions in pH enhance genome release, alter capsid dynamics and increase exposure of hydrophobic capsid residues at lower temperatures. The observation that the signals for exposure of genome and hydrophobic residues was reduced after pre-heating at elevated temperatures indicates that the changes that occur during heating are irreversible.

### Membrane interactions and pore formation

Having established that pH plays an important role in AiV uncoating, the effect of pH on the ability of AiV to permeabilise model membranes was investigated.

Purified AiV was incubated at a range of pH values with liposomes containing carboxyfluorescein (CF) dye at 37 °C. Florescent dye release was measured every minute for one hour. This revealed that AiV induces membrane permeabilisation in a pH dependent manner, with the rate of dye release increasing at lower pH values (Figure 3A). Dye release after one hour ranged from 12% at pH 7 to 90% at pH 4.9 (Figure 3B).

**Figure 3.**
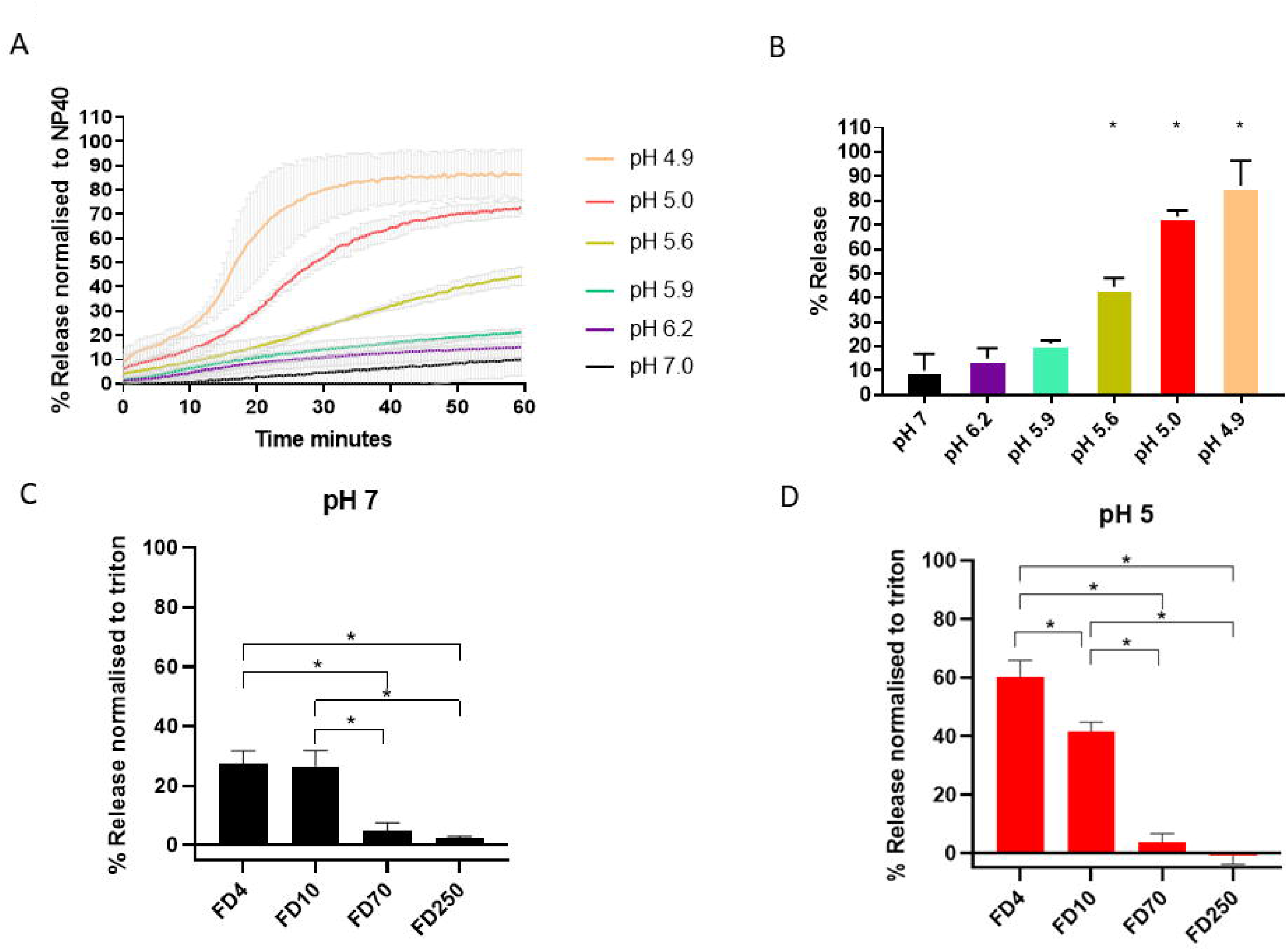
AiV-induced membrane permeability is enhanced by low pH. a) Permeability assay showing CF released from liposomes after mixing with AiV at pH 7.0, 6.2, 5.9, 5.6, 5.0 and 4.9. CF was detected by fluorescence measurements (excitation 492 nm/emission 512 nm) recorded every 30 seconds for 1 hour. b) End points of AiV induced membrane permeability. Data in panel a was normalised to the maximum signal induced by 0.1% NP40 at each pH value. c-d FD released after addition of 1 μg of purified AiV capsid liposomes containing FD of 4 kD (FD4), 10 kD (FD10), 70 kD (FD70) or 250 kD (FD250) at pH 7 (c) and pH 5 (d). Experiments were performed in triplicate and error is measured by s.e.m, * = p ≥ 0.01. Low pH is known to quench CF dye so all results are normalised to 100% release as determined by incubation by NP40 and 0% release as determined by liposome incubated alone. As low pH quenches CF/FITC fluorescence, the values were calculated as a percentage of full release by normalising to untreated liposomes (0% release) and 0.1% NP40 or triton (100% release).

In addition to increasing the rate of dye release, pH was also observed to change the profile of dye release curves. At pH 7.0, 6.2 and 5.9, AiV dye release curves were seen as a slope increasing at a steady constant rate, with the slope gradient increasing at the lower pH values. At pH 5.6, 5.0 and 4.9, there was an initial steady slope of release, before an exponential phase of release occurring at 20, 15 and 10 minutes receptively, before this levelled off (Figure 3 A). We have also investigated size selectivity using a dextran release assay. Samples of purified virus were incubated for one hour at pH 7.0 or pH 5.0 in the presence of liposomes containing FITC-labelled dextran of different sizes (4 kD (FD4), 10 kD (FD10), 70 kD (FD70) or 250 kD (FD250)) (Figure 3C-D). Release of dextrans was quantified by pelleting the liposomes and measuring the fluorescence in the supernatant. This revealed that AiV capsids preferentially released the two smallest dextrans (FD4 and FD10) and therefore appeared to form a size selective pore, consistent with previous published results with RV16 (2). The predicted size of the pore is consistent with the size necessary to allow passage of unfolded single stranded RNA (2). The effect of dye release for FD4 and FD10 was significantly higher at pH 5.0, giving further evidence that AiV induced pore formation is enhanced by low pH.

We have previously shown that for another picornavirus in which cell entry is dependent on endosome acidification, (RV16) the ability of the virus to permeabilise membranes was increased at lower pH values (2). In this previous study, the profile of RV16 dye release curves differed from AiV dye releases curves in the current study. For RV16 there was a high rate of dye release initially and the curve gradient gradually reduced over time, incubation at different pH values affected the rate of release but the profile remained the same. This could represent differences in uncoating dynamics of AiV and RV16.

### The role of VP0 in membrane permeability and pore formation in AiV

In other picornaviruses and picorna-like viruses, VP4 has been shown to be the component of the capsid that permeabilises membranes (1–7). For RV16, VP4 forms a size selective pore consistent with the size required for passage of single-stranded nucleic acid, specifically the first 45 amino acids are able to induce pore formation (7). Also residue 28 of PV has been shown to be involved with VP4 membrane permeability (5). In VP0 viruses such as AiV which do not undergo VP0 cleavage to form VP4 and VP2, it is hypothesised the N-terminus of VP0 will carry out this role. Given that the pore forming part of the enterovirus VP4 appears to be present in the first 45 amino acids, we investigated if there was conservation between VP4 sequences across the picornavirus family and if this was shared by VP0 viruses. Alignments of VP4 from a variety of different genera was performed using Muscle alignment (56). Alignments indicated a high degree of similarity of the amino acid properties of different picornavirus genera in the N-terminus of VP4, especially in the region of amino acids 20 and 35 (Figure 4A). This gives an indication that this region of VP4 may play a role in pore formation of all VP4 picornaviruses, consistent with this region containing the amino acid at position 28 mentioned earlier. We predict that this conserved motif is important for pore formation. We went on to look for conservation in VP0 viruses, comparing the first 109 amino acids of VP0 viruses from multiple genera using a Muscle alignment, this produced two groups of sequences which we refer to as ‘Kobu-like’ and ‘parecho-like’. The alignment shows that ‘parecho-like’ viruses lack strong conservation in this area and any other area of VP0. This is in contrast to ‘Kobu-like’ viruses which have a high degree of conservation of amino acid properties in the first 20 amino acids of VP0 (figure 4B). Strong conservation can be indicative of an important and essential function. Therefore this may suggest that the conserved VP4 motif in VP4 viruses and the conserved N-terminus of VP0 in ‘Kobu-like’ viruses, may have specific and essential functions. However, despite this conservation between VP4 sequences and between VP0 sequences, alignments between both VP4 and VP0 did not show obvious similarities (Data not shown). This suggests that there may be functional differences in how VP4 and VP0 interact with membranes.

**Figure 4.**
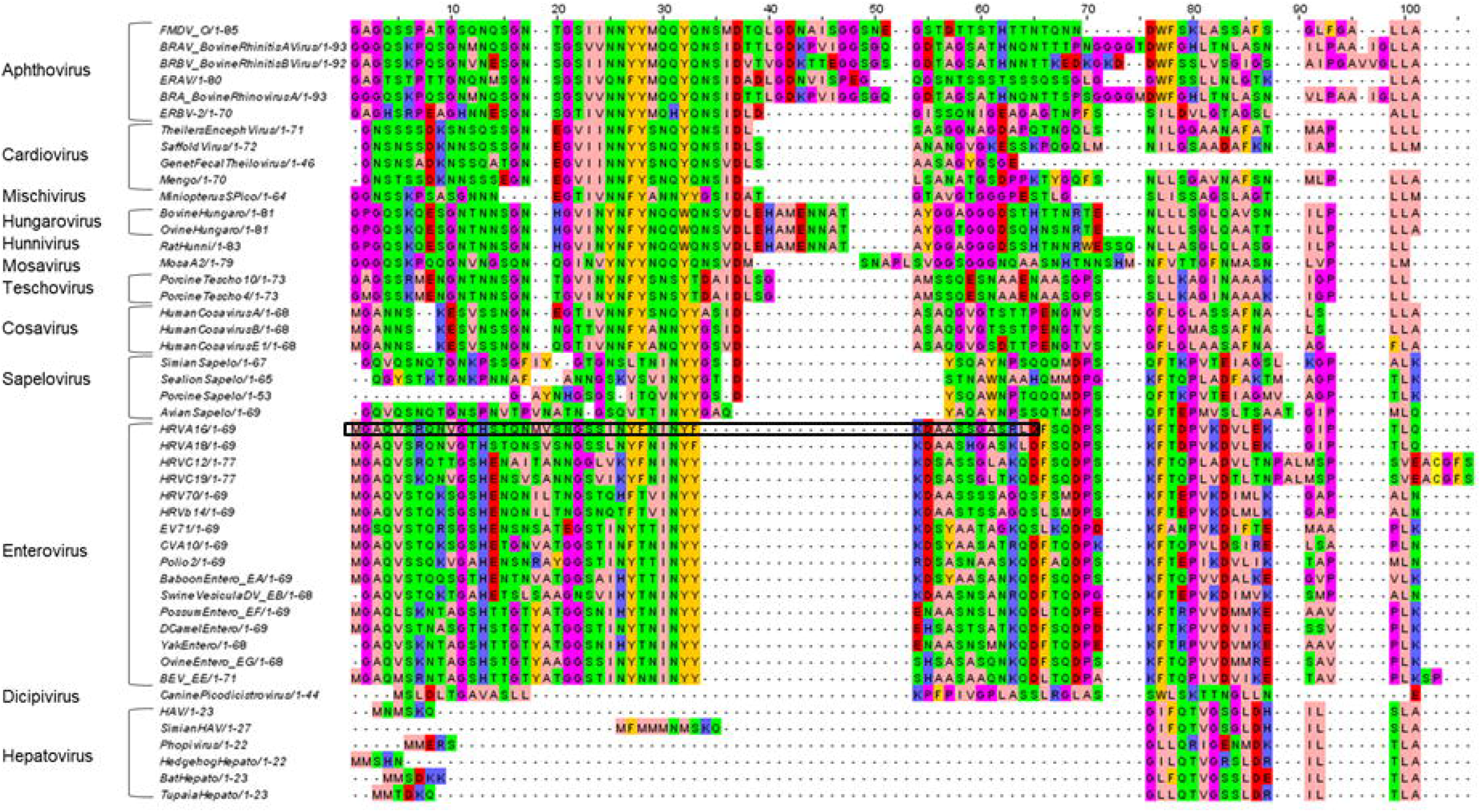
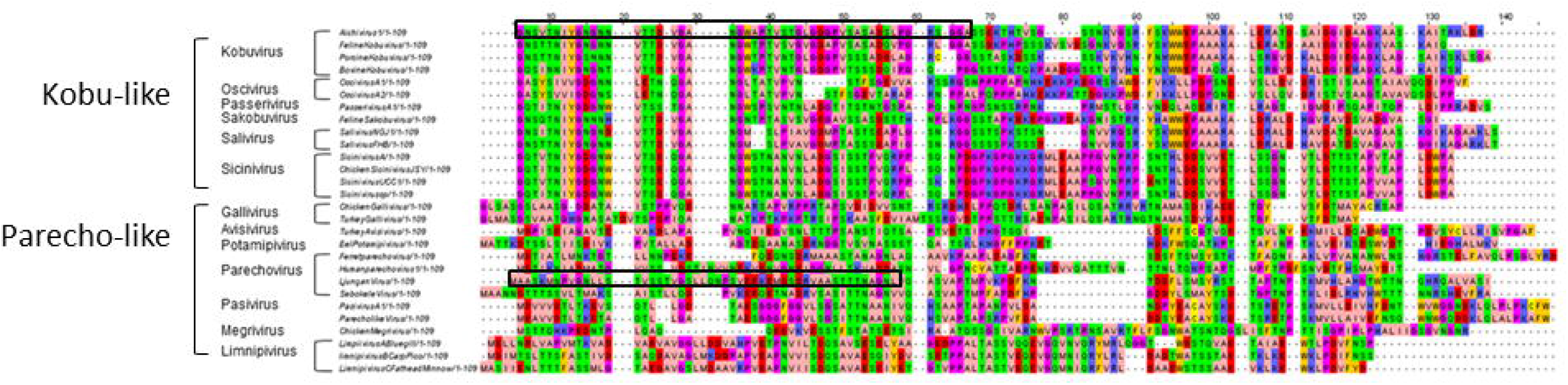
Alignments of VP4 and VP0 encoding picornavirus sequences. Alignments of picornavirus VP4 sequences (a) VP0 sequences (b) was carried out using the MUSCLE sequence alignment tool (56). The amino acids are coloured using the Zappo colour scheme as follows: aliphatic/ hydrophobic (pale pink), aromatic (orange), positively charged (purple), negatively charged (red), hydrophilic (green), conformationally special (magenta) and cysteine residues (yellow). Mention the new boxes. Peptide sequences regions for AiV, LV and RV are highlighted with a black box.

To test if the N-terminus of VP0 from other picornaviruses were able to induce pore formation, CF liposome assays were carried out using peptides from representatives of both groups, using the first 50 amino acids of AiV VP0 (AiV-VP0-N50) and the parechovirus, Ljungan virus (LV) VP0 (LV-VP0-N50) at pH 7. This revealed that AiV-VP0-N50 was able to induce dye release in a dose dependent manner, while LV-VP0-N50 was not (Figure 5 A-B). This is consistent with the VP0 sequences alignments where AiV and other ‘Kobu-like’ viruses show strong conservation in the N terminus of VP0, while LV and other ‘parecho-like’ viruses lack strong conservation in this area (Figure 5 A-B, Figure 4 B). The peptides used here were not myristoylated as it has previously been demonstrated that myristoylation is not essential for the replication of Kobuviruses and parechoviruses (28).

**Figure 5.**
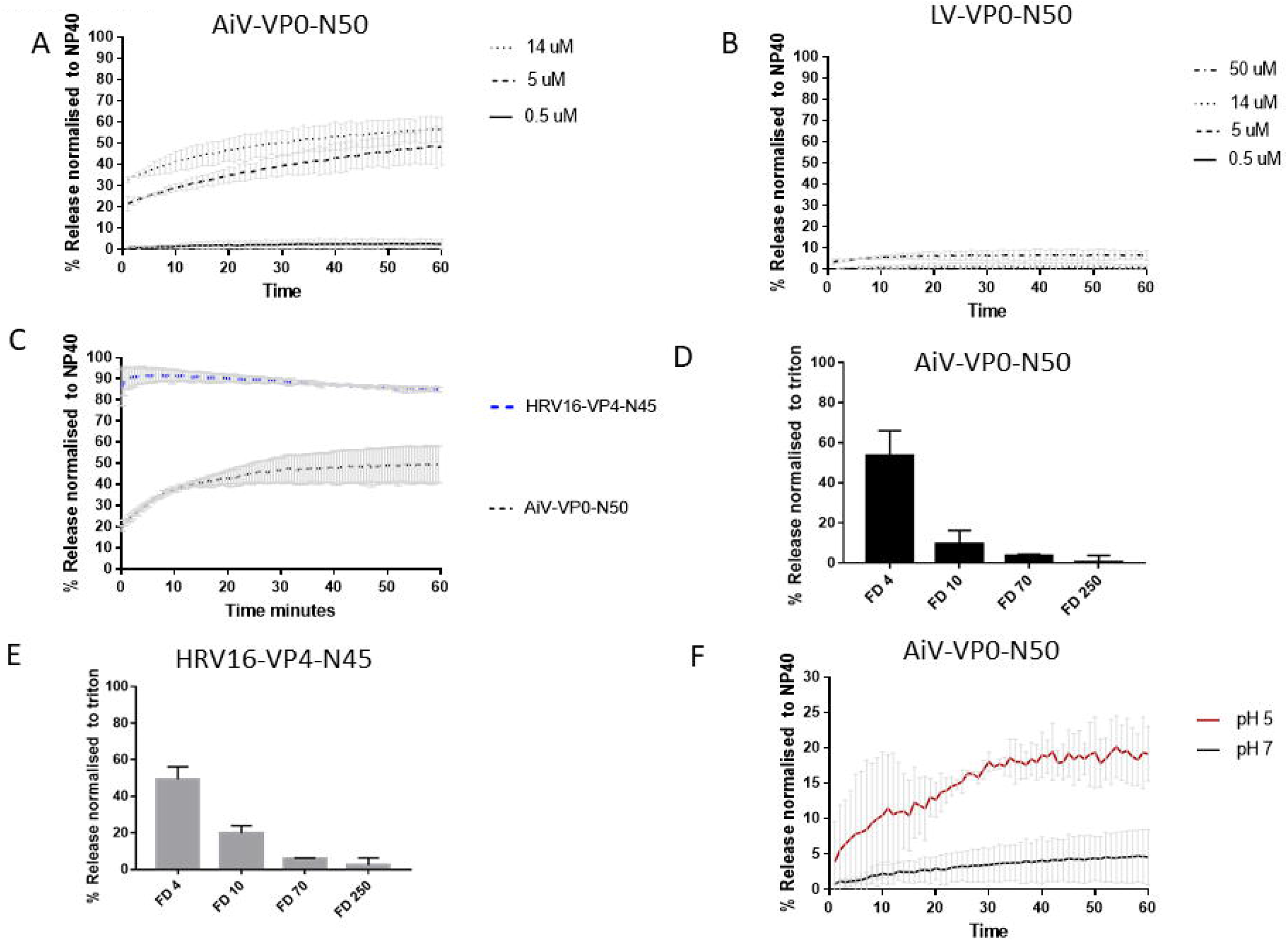
AiV VP0 N-terminal peptide is able to permeabilise membranes and form a size selective pore. Permeability assays showing CF or fluorescent dextrans (FD) released from liposomes after mixing with peptides. CF or FD was detected by fluorescence measurements (excitation 492 nm/emission 512 nm) and displayed as % of total release by detergent induced lysis. Assays were carried out at pH 7.0 unless otherwise stated. a) CF release over time after addition of peptide AiV-VP0-N50 at concentrations of 0.5 μM, 5 μM, 14 μM. b) CF release over time after addition of peptide ljungan virus (LV) LV-VP0-N50 at concentrations of 0.5 μM, 5 μM, 14 μM and 50 μM. c) CF release over time after addition of peptides AiV-VP0-N50 or RV16-VP4-N45 at concentrations of 5 μM. d) FD released after addition of 5 μM peptide AiV VP0 N50 to liposomes containing FD of 4 kD (FD4), 10 kD (FD10), 70 kD (FD70) or 250 kD (FD250). e) FD released after addition of 5 μM peptide RV16 VP4 N45 to liposomes containing FD of 4 kD (FD4), 10 kD (FD10), 70 kD (FD70) or 250 kD (FD250). f) CF release over time after addition of 0.5 μM peptide AiV-VP0-N50 at pH 5 or pH 7. All experiments were performed in triplicate and error is measured by s.e.m, * = p ≥ 0.01. As low pH quenches CF/FITC fluorescence the values were calculated as a percentage of full release by normalising to untreated liposomes (0% release) and 0.1% NP40 or triton (100% release). In FITC reactions pH of supernatant was neutralised by addition of a 2.5 M Tris pH 7.5 buffer. AiV-VP0-50N: GNSVTNIYGNGNNVTTDVGANGWAPTVSTGLGDGPVSASADSLPGRSGGA, LV-VP0-50N: MAASKMNPVGNLLSTVSSTVGSLLQNPSVEEKEMDSDRVAASTTTNAGNL, RV16-VP4-45N: MGAQVSRQNVGTHSTQNMVSNGSSINYFNINYFKDAASSGASRLD.

Although the AiV peptide was able to induce membrane permeability, this appeared to be at a lower level than previously observed for RV16 peptides (7). To test if there was indeed a difference in permeabilisation between RV16 and AiV peptides we performed the assays with the peptides in parallel. This revealed that AiV-VP0-N50 was less effective at inducing membrane permeability than an un-myristoylated RV16-VP4-N45 (Figure 5 C).

As it has been established that AiV-VP0-N50 is able to permeabilise membranes, its ability to form a size selective pore was compared with RV16 VP4 peptides, previously shown to form such a pore. To perform this AiV-VP0-N50 and RV16-VP4-N45 peptides were incubated for one hour at pH 7.0 in the presence of liposomes containing FITC-labelled dextrans of different sizes (4 kD (FD4), 10 kD (FD10), 70 kD (FD70) or 250 kD (FD250)) (Figure 5D-E). Release of dextrans was quantified by pelleting the liposomes and measuring the fluorescence in the supernatant. This revealed that, like RV16 VP4 and AiV capsids, AiV VP0 preferentially releases the smallest dextran FD4 and therefore forms a size selective pore (2).

Next the ability of AiV-VP0-N50 to permeabilise membranes was compared between pH 7.0 and pH 5.0. This revealed that pH 5.0 enhanced dye release of the peptide (Figure 5F). However, the dye release profiles at pH 5 differ between peptide and virus (Figure 5 F, Figure 3A). For the virus, a sharp increase followed by levelling off was observed, but for the peptide a constant gradual increase and no levelling was observed (Figure 5 C, Figure 3A).

## Discussion

In this study we sought to investigate uncoating and membrane interactions in the VP0-containing picornavirus AiV. This virus is of particular interest given the paucity of studies which have investigated uncoating in VP0-containing viruses. We have demonstrated that the N-terminus of AiV VP0 is a pore forming unit of the capsid, indicating its function is likely analogous/similar to VP4 in VP4-containing picornaviruses. Also using a combination of stability assays, membrane permeability assays and chemical inhibition we have demonstrated that pH plays a critical role in AiV uncoating and membrane interactions.

Here we have shown AiV requires endosome acidification for entry and using PaSTRy we demonstrated incubation of AiV at a pH of 5.6 and below enhances AiV genome exposure and alters the exposure of hydrophobic capsid proteins, this corresponds with the pH of late endosomes. The changes in PaSTRy assay profile were enhanced as pH was lowered even further, with a very dramatic shift occurring between pH 5.0 and 4.9. At present we are unable to explain this dramatic difference in profile, when the pH differs by just 0.1. At pH 4.9, but no other pH, an increase in hydrophobic protein signal is detected a few degrees before nucleic acid signal is detected. These events are thought to represent genome release and the externalisation of internal capsid proteins. After maximum nucleic acid signal is detected, there is a large drop in hydrophobic signal causing a trough. This may indicate that during uncoating, internal capsid proteins are externalised and then re-internalised after genome release. This would be consistent with structural data of AiV empty particles produced by heating capsids at neutral pH, which reveal that VP1 and VP0 are inside the capsid after genome release, indicating they may be externalised during uncoating and become reinternalised after genome release (50). However, PaSTRy assays performed on particles pre-heated (so that genome release had already occurred) no-longer showed this trough. This indicates the capsid rearrangements that occur during or after genome release are irreversible and if proteins are re-internalised after genome release their externalisation can no longer be initiated by heating. Furthermore, the biological relevance/implications of this *in vitro* observation still remains to be addressed. If re-arranged capsid proteins were inserted into a membrane first, it would likely be more difficult for them to be removed from the membrane and reinternalised into the capsid, as previous structural data (50), along with our biochemical data suggests they do in the absence of membranes.

Whatever the biological relevance though, our PaSTRy results and previous structures highlight a difference *in vitro* between AiV and enterovirus uncoating. Enterovirus structures show that VP4 is completely released from the capsid and the externalised termini of VP1 remains externalised after genome release (29, 35–37). Previously published data for enteroviruses with PaSTRy assay is consistent with this, with no apparent hydrophobic protein trough occurring after the nucleic acid peak (53, 54).

In addition to enhancing uncoating, incubation at a more acidic pH also enhances the ability of AiV to permeabilise model membranes. When AiV is incubated at pH 7.0 it induces relatively low levels of dye release, reducing the pH to 6.2 and 5.9 caused a moderate increase in dye release. In PaSTRy assays these pH values did not affect genome exposure and had similar hydrophobic protein profiles to pH 7.0. When the pH is lowered to 5.6 and below, the rate of AiV induced dye release increases significantly before levelling off, this affect becomes more pronounced as the pH decreases further (Figure 3). These significant increases in dye release coincide with PaSTRy assay profiles with enhanced hydrophobic signal. This suggests that externalisation of hydrophobic capsid protein residues enhances AiV induced membrane permeabilisation, this is consistent with models of enterovirus induced membrane permeabilisation (45). The liposome assay profiles observed for AiV differs from what we have seen previously for the enterovirus RV16. For RV16, particles were able to induce dye release at neutral pH, dye release was enhanced at low pH but it increased the rate of release gradually, rather than a sudden increase in release and then levelling off which we observed for AiV (2). This may indicate that low pH plays a more critical role in AiV particle alterations and membrane interactions than for RV16.

Given that incubation of AiV at low pH increases the level of hydrophobic protein residues detected in PaSTRy and enhances membrane interactions, it is likely that low pH induces externalisation of capsid components essential for AiV membrane interactions. It might therefore be expected that free peptide would induce higher levels of membrane permeability at physiological pH than virus. However, this was not observed, VP0-N-50 peptide at 0.5 μM induced lower amounts of dye release than virus containing the equivalent of 0.1 μM of VP0-N-50 at pH 7.0 and pH 5.0. As free peptide is less efficient at inducing membrane permeability than the virus, this indicates that either additional components of the capsid are also involved in membrane permeabilisation or that VP0 N must be physically attached to the capsid to maintain its optimal pore forming conformation. Low pH also enhances the ability of the VP0-N-50 peptide to form a pore, this is not surprising given that this would be the natural environment that it would be required to form a pore in. This is consistent with observations that the ability of RV16 VP4 protein induce pore formation is enhanced at acidic pH (2).

Furthermore, using an N-terminal peptide of AiV VP0, we demonstrated VP0 forms a size-selective pore in model membranes consistent with a pore size able to release a molecule of single stranded RNA. This demonstrates the N-terminus of AiV VP0 plays an analogous function to VP4 of other picornaviruses in terms of membrane permeabilisation (1–6). However, the N-terminus of VP0 does not appear to be the pore-forming component of all VP0-containing picornaviruses, as an N-terminal peptide for LV, did not induce dye release from liposomes.

Sequence comparison of VP0 N-termini reveal that the VP0-N termini of AiV and other ‘Kobu-like’ VP0 viruses are well conserved, this suggests that the N-terminus of VP0 likely plays a pore forming role for all ‘Kobu-like’ VP0 viruses. Similar levels of homology are seen between VP4 sequences in other picornavirus genera. However, comparison of LV and other ‘parecho-like’ VP0 viruses revealed that VP0 N-termini are not well conserved, this would be consistent with it not being involved in membrane interactions in these viruses. Taken together, this seems to indicate that the N-terminus of the VP0 of viruses in ‘Kobu-like’ viruses possess the ability to form a pore, whilst the ‘parecho-like’ group lack N-terminal membrane permeabilisation activity.

The ability of other AiV proteins to interact with membranes is yet to be determined. For enteroviruses, it has been demonstrated that the N-terminus of VP1 is essential for attachment to model membranes in flotation assays. The N-terminus of VP1 is internal in native particles and is released during conversion to A-particles. It was shown that in flotation assays, A-particles bound to model membranes, while native particles and A-Particles where VP1 N-terminus had been cleaved by proteolytic digestion were unable to bind model membranes (30). Furthermore, when A-particles bound to model membranes were subjected to proteolytic digestion, the N-terminus of VP1 remained within the membranes (30). Further demonstrating that VP1 is required for membrane attachment in enteroviruses (30). If consistent with enteroviruses, the AiV N-terminus of VP1 will become externalised and be involved in attachment to the endosomal membrane. However, unlike in VP4-containing-picornaviruses, AiV VP0 remains attached to the capsid, so it is possible that it may play an important role in attachment alongside or instead of VP1.

### Conclusion

This study is the first to characterise effects of pH on the uncoating and membrane interactions of a VP0-containing picornavirus. We have shown that the N-terminus of VP0 can play a role in pore formation but not in all VP0-contianing-picornaviruses. We have also demonstrated that AiV behaves differently in functional uncoating assays compared to enteroviruses. Together with previous structural studies this indicates that AiV capsids likely undergo different structural changes in the capsid to initiate membrane interactions and uncoating than VP4-containing-picornaviruses. AiV also appears to have a greater dependence on pH to facilitate externalisation of membrane interacting components than enteroviruses. Further characterisation will be required to determine the exact uncoating mechanism of AiV.

## Materials and Methods

### Cell lines and virus

Vero cells were obtained from the Central Services Unit at The Pirbright institute and propagated in DMEM containing 10% fetal bovine serum (FBS) and 50 μg/ml penicillin and streptomycin at 37° C in a humidified atmosphere containing 5% CO_2_. AiV strain A846/88 (GenBank no. BAA31356.1) was obtained from Prof David Stuart and Dr Elizabeth Fry at the University of Oxford. Virus was propagated by inoculating Vero cells at MOI 1 and incubating at 37 °C in a humidified atmosphere containing 5% CO_2_ for 24 hours before the supernatant was harvested.

### Peptides

Peptides were synthesised by Peptide Protein Research Ltd using the PeptideSynthetics service. The sequences were; AiV-VP0-50N: GNSVTNIYGNGNNVTTDVGANGWAPTVSTGLGDGPVSASADSLPGRSGGA LV-VP0-50N: MAASKMNPVGNLLSTVSSTVGSLLQNPSVEEKEMDSDRVAASTTTNAGNL RV16-VP4-45N: MGAQVSRQNVGTHSTQNMVSNGSSINYFNINYFKDAASSGASRLD

### Chemical inhibition

Vero cells were seeded into a 6-well plate at 3×10^5^ cells in 2 ml of 10 % FBS-DMEM per well and incubated at 37 °C overnight. Medium was removed from the wells and the cells were pre-treated with media containing NH_4_Cl (ammonium chloride) (Sigma-Aldrich) (2, 10, 20, 40 mM), for 2 hours at 37 °C. Cells were then incubated on ice with virus to allow attachment (MOI = 1) for 30 minutes in the presence of inhibitor. Unbound virus were removed, and cells were washed with PBS, before adding 2 ml of warm serum-free DMEM containing inhibitor to the wells. For time of addition studies cells were infected with AiV at MOI 1 and media was replaced with 40 mM of NH_4_Cl at with 0, 1, 2, 3 or 4 hours post infection. After incubation overnight at 37 °C for 20 hours the supernatants were harvested and the virus titres determined by plaque assay.

### Plaque assay

Six well plates were seeded with 3×10^5^ Vero cells per well. The following day AiV samples to be titrated was serially diluted in serum-containing medium. Media was removed from wells and 200 μl of each serial dilution were added to individual wells and incubated for 2 hours. After 2 hours, supernatant was removed and 2 ml of serum-containing medium with 1% agarose at 42 °C was added to each well and allowed to solidify. Plates were incubated at 37 °C in a humidified atmosphere containing 5% CO_2_ for 72 hours. Monolayers were fixed and stained with 1 ml of 4% formaldehyde, 1% crystal violet, 20 % ethanol in PBS, plaques counted and titre expressed as PFU/ml of starting material.

### Purification of virus

Infected cell cultures were lysed by addition of NP40 to make the solution a final concentration of 0.5% NP40 and freeze-thawing three times. Lysates were incubated for 3 hours at 37 °C in the presence of DNAse (10 μM) and clarified by centrifugation. Clarified supernatants were concentrated by precipitation with 8% PEG 8000 overnight and centrifugation at 100,000 RCF for 1 hour. The resulting pellet was resuspended in PBS, pelleted through a 2 ml cushion of 30% (w/v) sucrose in PBS at 125,755 RCF for 2 hours, resuspended in PBS and subjected to sedimentation in a sucrose density gradient comprising 15–45% (w/v) sucrose in PBS at 237,000 RCF for 50 minutes. Sucrose gradients were fractionated and purified virus was quantified by absorbance at 260 nm. Sucrose was removed using a Zeba column (ThermoFisher Scientific) following the manufacturer’s instructions.

### Particle Stability Thermal Release Assay (PaSTRy)

Virus particle alterations were characterized by a thermofluorometric dual dye-binding assay using the nucleic acid dye SYTO9 and the protein-binding dye SYPRO orange (both from Invitrogen). Reaction mixtures of 50 μl containing 1.0 μg of purified virus, and 0.1M Citric acid 0.2M Sodium phosphate dibasic buffer at either pH 7, 6.2, 5.9, 5.6, 5.0 or 4.9 were mixed and incubated at either room temperature, 56 °C, or 59 °C for 10 minutes and then chilled on ice for 2 minutes. Reaction mixes were then made to 5 μM SYTO9, 150X SYPRO orange, and ramped from 25 to 95°C, with fluorescence reads taken at 1°C intervals every 30 s within the Stratagene MX3005p quantitative-PCR (qPCR) system.

### Preparation of liposomes

Liposomes comprising of phosphatidic acid, phospatidylcholine, cholesterol and rhodamine-labelled phosphatidylethanolamine (Avanti Polar Lipids) (molar ratios 44.5:44.5:10:1 respectively) were prepared as previously described (6) by rehydration of dried lipid films in 107 mM NaCl, 10 mM Hepes pH 7.5 and extrusion through 400 nm pore-size membranes using a mini-extruder (Avanti Polar Lipids). The lipid concentration of liposome preparations was estimated by comparing the level of rhodamine fluorescence in the liposome sample relative to samples of rehydrated lipids of known concentration. The expected diameter (average 400 nm) and size distribution of liposomes was confirmed by dynamic light scattering (Malvern Zetasizer μV). Liposomes containing carboxy-fluorescein (CF) (Sigma) were prepared by rehydrating lipids in the presence of 50 mM CF, 10 mM Hepes pH 7.5. Liposomes containing FITC-conjugated dextrans (FD; Sigma) were prepared by rehydrating lipids in the presence of 25 mg/ml FD, 107 mM NaCl, 10 mM Hepes pH 7.5. Liposomes containing CF or FD were purified from external fluorescence by multiple cycles of ultracentrifugation (1) and resuspended in 107 mM NaCl, 10 mM Hepes pH 7.0 or 0.1M Citric acid 0.2M Sodium phosphate pH 7, 6.2, 5.9, 5.6, 5.0 or 4.9.

### Membrane permeability assays

Membrane permeability was measured by detecting the release of fluorescent material from within liposomes. Purified liposomes containing CF or FD were added to test substances (peptide, virus or mock controls in typical volume 5 μl) to give typical final concentrations of 50 μM lipid, 107 mM NaCl, 10 mM HEPES pH 7.4 or 0.1M Citric acid 0.2M Sodium phosphate pH 7.0, 6.2, 5.9, 5.6, 5.0 or 4.9 and total volume of 100 μl. Reagents and plastic-ware were pre-equilibrated to the reaction temperature (25°C or 37°C).

For CF release, reactions were assembled in 96-well plates and membrane permeability detected in real time by the release, dequenching and increase in fluorescence of CF. Measurements were recorded every 30 s for 1 hr using a fluorescence plate reader with excitation and emission wavelengths of 485 nm and 520 nm respectively (Plate CHAMELEON V, Hidex). Initial rates were calculated from the linear slope of lines generated from the initial four data points.

For experiments investigating the effect of pH on the induction of membrane permeability, CF release reactions were assembled with 0.1M Citric acid 0.2M Sodium phosphate pH 7.0, 6.2, 5.9, 5.6, 5.0 (instead of HEPES). CF fluorescence is quenched at low pH. As low pH quenches CF fluorescence the values were calculated as a percentage of full release by normalising to untreated liposomes (0% release) and 0.1% NP40 (100% release).

FD release reactions were assembled with 10 mM citric acid and 10 mM sodium phosphate at pH 5 and pH 7. Reactions were incubated for 1 hr, liposomes pelleted at 100,000× g for 30 mins and pH of supernatant was neutralised by addition of a 2.5 M Tris pH 7.5 buffer. Fluorescent signal in the supernatant was measured using the plate reader as above. The signal pelleted liposomes signal was then released by the addition triton to calculate a 100% release signal.

## Acknowledgments

We would like to thank Prof David Stuart and Dr Elizabeth Fry at University of Oxford for the kind gift of AiV.

